# An efficient pipeline for creating metagenomic-assembled genomes from ancient oral microbiomes

**DOI:** 10.1101/2024.09.18.613623

**Authors:** Francesca J. Standeven, Gwyn Dahlquist-Axe, Camilla F. Speller, Conor J. Meehan, Andrew Tedder

**Affiliations:** School of Chemistry and Biosciences, University of Bradford; Department of Anthropology, University of British Columbia; Department of Biosciences, Nottingham Trent University

## Abstract

Metagenomic-assembled genomes (MAGs) are difficult to recover from ancient DNA (aDNA) due to substantial fragmentation, degradation, and multi-source contamination. These complexities associated with aDNA raise concerns about whether bioinformatic tools intended for interpreting modern DNA are suitable for reconstructing ancient MAGs. Using simulated modern and ancient data, we investigated: 1) how using binning tools designed for modern DNA affects our ability to effectively construct MAGs from ancient genomes; 2) the performance of three different binning tools for aDNA samples; and 3) whether a ‘one size fits all’ approach is suitable for ancient metagenomics. We established that binning tools for modern DNA performed efficiently on simulated modern and ancient DNA. When applied to ‘real’ archaeological DNA spanning 5000 years, we retrieve high-confidence MAGs in most cases.

## Introduction

Recovering meaningful genomic data at the operational taxonomic unit (OTU) level from ancient microbiome (microbial community) samples requires the use of metagenomic-assembled genomes (MAGs). This differs from a single ‘species’ approach, in that a MAG represents a microbial genome using a group of sequences from assembled genomes with similar traits (Yang et al. 2021) rather than identifying species based on individual sequences. Reconstructing MAGs, therefore, offers us significant insight into uncultured microbial genomic diversity (Churcheward et al. 2022), can enable the detection of new species and offer evidence into their potential functions in a dynamic ecosystem (Yang et al. 2021), and the integration of ancient MAGs enriches this analysis. For example, a recent study by Klapper et al. (2023) reconstructed ancient Chlorobium MAGs which uncovered unmodified metabolites, allowing for the observation of functional traits over millennia.

Archaeological science has delved into the historical events and lifestyle transitions that had a huge impact on the health of past societies, and the creation of ancient oral MAGs has allowed for an understanding of the origins and evolution of microbial species and strains in the oral cavity. For example, Wagner et al. (2014) reassembled a *Yersinia pestis* draft genome from dental calculus, revealing a novel branch dating back to the Justinian plague. This study showed how their strain differs from strains connected with future pandemics caused by *Y. pestis*, such as the Black Death. Other research has also used *Y. pestis* genomes to evidence its establishment during the Bronze Age (Spyrou et al. 2018). In another study, the reconstruction of *Methanobrevibacter* genomes from historical dental calculus across different regions has revealed previously unknown species that demonstrate a decline of the genus over thousands of years due to changing diets (Granehäll et al. 2021). Metagenomic assembled *Olsenella* sp. Oral taxon 807 genomes showed several missing components related to virulence and biofilm formation compared to modern genomes, implying that the species has recently evolved and adapted to thrive in the oral microbiome, most likely due to stronger selective pressure exerted by dietary changes (Quagliariello et al. 2022).

Due to the poor quality and quantity of aDNA, combined with high contamination levels (Gaeta 2021), ancient metagenomics is still an extremely challenging field of study; therefore, the efficiency of modern MAG generation techniques when applied to aDNA remains unclear. Depurination, a primary mechanism for miscoding lesions in aDNA, converts cytosine into uracil which is often mistaken for thymine (Dabney et al. 2013). The depurination process can also result in highly fragmented DNA (Kistler et al. 2017). Hydrolytic depurination, followed by beta-elimination, causes single-strand breaks in DNA when exposed to acidic environmental conditions (Dabney et al. 2013). The extent of this damage depends on the location and duration of the DNA’s burial; nevertheless, aDNA fragments are generally very short ranging from 40–500 bp (Dabney et al. 2013). The chemical processes that cause damage have the potential to influence the performance of modern MAG assembly methods on aDNA. aDNA is also prone to environmental and modern contamination because it can be situated in the ground for thousands of years, making it susceptible to contamination from environmental microbes in the soil (Kazarina et al. 2019). Likewise, the degraded nature of aDNA means that it is frequently low template, and thus at more risk from modern contamination resulting from post-excavation handling including the laboratory environment (Llamas et al. 2017). Overall, both damage and contamination affect the ability to create ancient MAGs with the same degree of confidence as modern DNA because the effects of these traits on metagenomic assembly are not fully understood.

Although the field of aDNA is quickly progressing, it is still difficult to retrieve high sequencing coverage for historical samples with short read lengths and various challenges can arise when used with bioinformatic software packages designed for modern long-read DNA. Commonly applied data pre-processing approaches exacerbate these issues by further shortening the read length and likely impacting MAG assembly. For example, necessary decontamination procedures such as BBduk ( Joint Genome Institute 2023) have the potential to greatly reduce read length by removing vast amounts of possibly contaminated sequences and can cause short, uneven counts in paired-end reads. Additional tools, such as Trimmomatic (Bolger et al. 2014) or PEAR (Zhang et al. 2014), must sometimes be applied to repair these files by evenly cutting off base pairs that result in even more sequence reduction. Therefore, the short read length in ancient metagenomics emphasises the need for experimentation in genome assembly techniques.

MAGs are valuable resources for archaeological scientists and microbiologists. Therefore, utilising efficient and effective assembly methodologies is paramount for ensuring accurate and authentic ancient genomes. Reconstructing MAGs involves genomic binning where organised contigs (contiguous fragments) are sorted into groups of high similarity, or ‘bins’, whereby each bin corresponds to a MAG (Setubal 2021). A bin is an individual genome reconstruction made from contigs that are similar in composition. Binning methodologies vary, but a common theme involves comparing contigs to reference genomes as part of the decision tree. Several successful binning tools are available for modern DNA, such as Maxbin (Wu et al. 2016), CONCOCT (Alneberg et al. 2014), and MetaBAT (Kang et al. 2015), as well as many MAGs pipelines that have one or more binning tools already implemented (Uritskiy et al. 2018). It has been argued that a combination of binning tools can strengthen binning by the application of many independent variables for contig clustering (Hug 2018). After applying various binning tools, overlaps and redundancy can be determined. For example, the programme DAS Tool can be used to combine various binning algorithms to rewrite and calculate an optimised, non-redundant selection of bins from a single assembly (Sieber *et al*. 2018). In light of all of the issues with aDNA, it is unclear to what extent existing binning methods and pipelines are effective, and whether further optimisation is required. Despite recent pipeline developments with functionalities geared towards ancient MAG production (Krakau et al. 2022), systematic comparisons of modern MAG binning tools on ancient DNA are lacking.

MAGs give a degree of reliability because they require numerous reads to form their assembly and we can calculate their completeness and contamination; however, the performance of modern DNA binning techniques on ancient sequences is understudied and could result in unreliable MAG outcomes, such as false positives and false negatives. To determine whether standard tools for modern MAG creation are suitable for aDNA, we tested our binning pipeline on simulated ‘modern’ and ‘ancient’ data by quantifying MAG success and binning performance. Our method was then applied to a real aDNA dataset obtained from archaeological dental calculus to understand if binning tools developed for modern DNA have aDNA limitations and if factors such as short read length is a barrier to MAG recovery success in aDNA.

## Materials

### Simulated data

To disentangle the impact of aDNA deamination (damage) on the process of MAG recovery and reconstruction, simulated metagenomic populations were created using InSilicoSeq v1.6.0 (Gourlé et al. 2019). Twenty modern populations were simulated to reflect paired-end (PE) 300 bp MiSeq reads. ‘Populations’ comprised three replicate samples, each of which contained reads taken from 10 microbial genomes (not limited to oral microbial genomes) pulled randomly from NCBI. The frequency of each OTU (per sample) was determined at random (Gourlé et al. 2019). Simulated modern samples then underwent simulated delamination using ‘deamSim’ from gargammel v1.1.4 (Renaud et al. 2017) with the following default parameters -damage 0.03, 0.4, 0.01, 0.7 to damage nucleotide bases. Fragmentation is not explicitly modelled in this dataset. Information on species and abundance for the simulated datasets is provided in Table S1.

### Archaeological data

The ancient oral microbiome samples used in this research were selected from data reported by (Standeven et al. 2024). The dataset comprises 30 pre-industrial (ca. 2,300 BC–1750), 21 Industrial (ca. 1750–1850), and 54 post-industrial (ca. 1837–1901) samples. Sample ID, European Nucleotide Archive (ENA) accession number, sample type, site location, historical period, and read length of samples are listed in Table S2.

## Methodology

Our simulated data and real archaeological samples were run through our hybrid-scripted MAGs pipeline (**Fig 1**) described below. The pipeline was first applied to the simulated data to test its performance before real archaeological samples. Then, real archaeological samples were run through the pipeline via period batch (see materials) and then per sample to comprehend how ancient characteristics such as low read count affect binning. Damage authentication and decontamination protocols were only applied to real archaeological data with matched laboratory controls.

**Figure 1.**
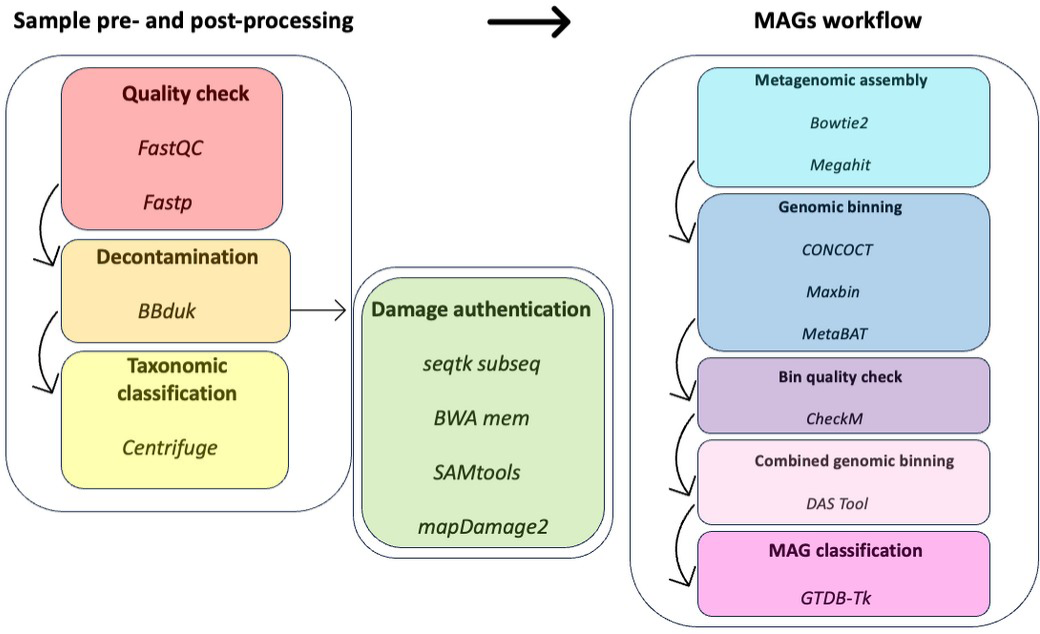
MAGs assembly and QC workflow.

### Hybrid-scripted MAGs pipeline

#### Sample pre-processing

Standeven *et al*. (2024) provide a full overview of quality control, decontamination, and damage authentication protocols for ancient metagenomic sequences. FastQC v0.11.9 (Babraham Bioinformatics 2019) was used with default parameters to quality-check samples, pre-and post-filtering. Fastp v0.23.2 (Chen et al. 2018) was used with default parameters to perform read filtering, trimming, adapter removal, and mismatched bp repair on raw metagenomic sequencing data. To minimise the impact of laboratory and environmental contamination, BBduk ( Joint Genome Institute 2023) was used with default parameters to identify and remove homologous sequences found in laboratory and environmental controls - when matching environmental samples (bone and tooth root) were available (Standeven et al. 2024) - from raw trimmed reads. Centrifuge v1.0.3 (Kim et al. 2016) was performed with default parameters to screen for OTUs. Non-human reads were retained from Centrifuge outputs and converted into post-Centrifuge fastq files using seqtk v1.4 ‘subseq’ (Li 2024). To authenticate ancient sequences, we retained only human reads from Centrifuge outputs and converted them into fastq files using seqtk ‘subseq’ which were then mapped to the human genome (hg38) (NCBI 2013) using BWA mem v0.7.17. SAMtools v1.12 (Danecek et al. 2021) was used to prepare the files for mapDamage2 v2.2.2 ( Jónsson et al. 2013) which was run with default parameters.

#### Metagenomic assembly

Metagenomic assembly of post-Centrifuge fastq files was performed using Megahit v1.2.9 (Li et al. 2015), a tool used to assemble large and complex metagenomics *de novo* datasets. The parameters --kmin-1pass --no-mercy were applied for memory efficiency while recovering low-coverage sequences. To ensure file compatibility with later tools, contigs produced by Megahit were reformatted into fasta files using the microbial ‘omics platform anvio v7 (Eren et al. 2020). Anvio was used to filter out short contigs <2500bp (-l 2500, --simplify-names) to promote good MAG quality. Studies recommend using contigs no less than 1000 bp for binning (Meziti et al. 2021) and given the ancient nature of our samples, we aimed to be especially cautious and apply stringent parameters to minimize the risk of false positives. Bowtie2 v2.5.0 (Langmead and Salzberg 2012) was used to map raw reads back to the assembly with the parameter -no-unal to suppress unaligned reads.

#### Genomic binning

Contigs were binned using three individual binning tools: MetaBAT v2.12.1 (Kang *et al*. 2015) with parameters -m 2000 for minimum contig size, and -- saveCls to save cluster memberships; CONCOCT v1.1.0 (Alneberg *et al*. 2014) with parameters -c 10000 for maximal number of clusters and a default minimum contig length threshold of 1000; and Maxbin2 v2.2.7 (Wu *et al*. 2016) via anvio with a default minimum contig length threshold of 1000.

To obtain the best quality MAGs, DAS Tool v1.1.6 (Sieber *et al*. 2018) was used with default parameters to merge, de-replicate, and select good-quality bins (≥75% completeness) from each binning tool. After DAS Tool selected bins, bins were then designated for further classification based on a completeness threshold of ≥75%.

#### Quality checking

We utilised CheckM v1.2.2 (Parks *et al*. 2015) with the automatic lineage workflow (lineage_wf) to assess percentages of completeness and contamination in MAGs. We conformed to recent guidelines (Bowers *et al*. 2017) that group MAGs into either ‘low-quality draft’ (<50% complete with <10% contamination), ‘medium-quality draft’ (≥50% complete with <10% contamination), or ‘high-quality draft’ (>90% complete with <5% contamination) genomes.

#### MAG taxonomic classification

To assign taxonomic labels to the MAGs, we used the Genome Taxonomy Database Toolkit v1.0.2 (GTDB-Tk; Chaumeil *et al*. 2020) using the workflow ‘classify_wf’ to process genomes using both bacterial and archaeal markers.

### Quantifying MAG success

After running the simulated data through the pipeline shown in **Fig 1**, MAGs success was measured in the following way: Firstly, to determine if MAG classifications corresponded to the correct OTUs, FastANI v1.33 ( Jain et al. 2018) was used to calculate the average nucleotide identity (ANI) between MAG files and original OTUs (see Table S3 for FastANI outputs detailing query and reference genome, ANI, fragment mappings and total query fragments). Then, a Python script was run to query GenBank via Entrez (the NCBI retrieval system) for tax ID numbers for each initial OTU and each MAG in FastANI files, then the tax IDs were matched to calculate true and false positives (see Table S4 for information on tax ID and tax name corresponding with OTUs). A paired t-test was performed on the number of true and false positive MAGs between modern and ancient sample sets to measure overall MAG retrieval success.

It is important to understand factors that may influence failure to retrieve MAGs for each OTU in a given sample. To disentangle factors linked to MAG retrieval success, such as OTU abundance in the initial sample, OTU genome size, and OTU read count, we used FastANI v1.33 ( Jain et al. 2018) to attribute MAGs from both modern and ancient simulated samples to an OTU. We used an ANI of 96% (Ciufo et al. 2018) to determine MAG identity. MAG retrieval success (MAG produced or not) was then compared to OTU abundance in the initial sample population pool and OTU genome size, as well as OTU abundance and read count.

### Quantifying binning tool performance

To determine whether binning using a single binning tool performs adequately for aDNA, or whether a multi-binning tool approach is required, we compared the frequency in which all three binning tools produce MAGs for the same OTU above our acceptance threshold of 75% completeness, before the DAS tool step. This analysis was performed on simulated ancient data.

### Applying modern genome assembly methods to real archaeological data

After running the pipeline on simulated data, the MAG pipeline described above (**Fig 1**) was applied to real archaeological dental calculus to understand if binning is effective on single samples in comparison to groups of samples separated by historical period.

## Results

### MAG success on simulated data

85.5% (171 out of 200) of simulated modern OTUs and 79.5% (159 out of 200) of simulated ancient OTUs were represented by MAGs. The paired t-test indicated no significant difference (p-value = 0.94) in MAG success between modern and ancient simulated groups. Neither simulated group had evidence of false positive MAGs. While MAG retrieval success was not significantly different between simulated modern and ancient samples, the MAG pipeline failed to retrieve MAGs (false negatives) in simulated ancient samples with a higher frequency (20.5% in ancient vs. 14.5% in modern). As expected, low OTU abundance, shown with varying read count (sample index), in both simulated sets results in failure to retrieve MAGs (**Fig 2a & c**), although we also see OTUs with high abundance (>10%) fail in both sequence types. This appears to correlate with OTU genome size (**Fig 2b** & **d**), with larger genomes increasing the likelihood of failure to retrieve MAGs even with moderate to high abundance.

**Figure 2.**
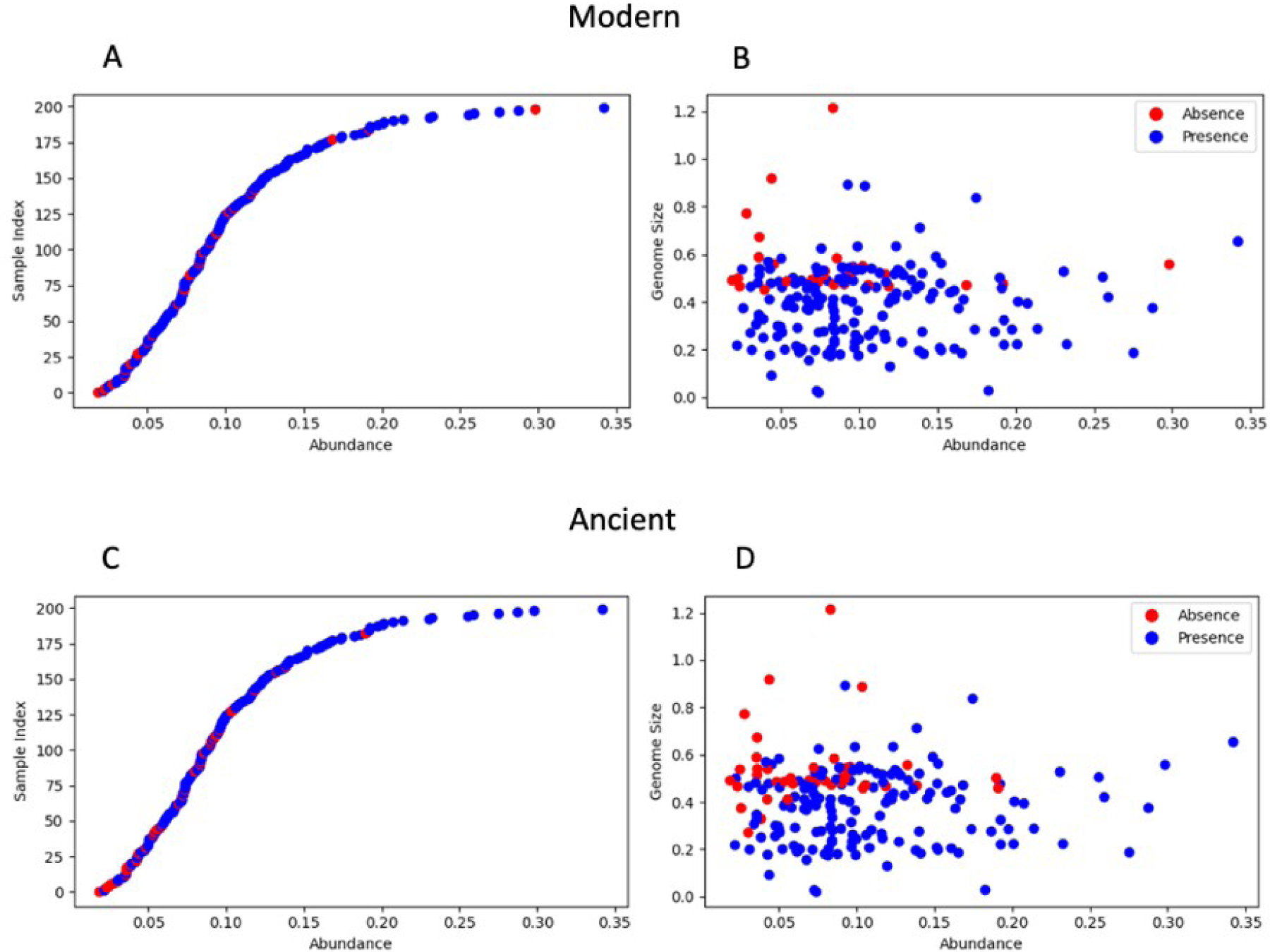
MAGs presence (blue) and absence (red) based on sorted abundance and read count (a & c) and abundance and genome size (b & d) for both simulated modern and ancient groups.

### Binning tool performance on simulated data

Binning tools all achieve overlapping bins in only 44% of cases. In the remaining 56% of cases where one or more of the binning tools retrieved a unique bin not retrieved by the other tools, each binning tool (CONCOCT = 35%; Maxbin = 40%; and MetaBAT = 25%) retrieved a proportionately similar number unique bins, although CONCOCT and Maxbin performed slightly better.

### Archaeological MAGs

Following the simulated data experiments, we ran the pipeline on our real archaeological metagenomic dataset to examine binning performance on samples via period and per sample.

With one exception, none of the binning tools were able to retrieve MAGs from single samples. A medieval tooth sample (LC26T; 125 bp) was the only instance in which MAGs (with varying quality) were identified. While some of these MAGs corresponded with those found in the pooled samples from the pre-industrial period, there was not a complete overlap. MAG retrieval was much more successful when samples were grouped into meaningful period categories. CheckM completeness and contamination levels for all bins (before DAS Tool selection) across all periods are summarised below and presented in **Table 2** and **Fig 3**. A more detailed summary of information on the final MAGs, including completeness and contamination, GTDB classification, and genome quality can be found in Table S5. In total, 13 pre-industrial bins (CONCOCT=6, Maxbin=2, MetaBAT=5), 7 Industrial bins (CONCOCT=4, Maxbin=0, MetaBAT=3), and 18 Post-Industrial bins (CONCOCT=7, Maxbin=2, MetaBAT=9) from the DAS Tool bin list output met the ≥75% completeness threshold. CheckM values indicate that the majority of bins (34 out of 38 bins) range from medium (≥50% complete with <10% contamination) to high (≥90% complete with <5% contamination) quality.

**Figure 3.**
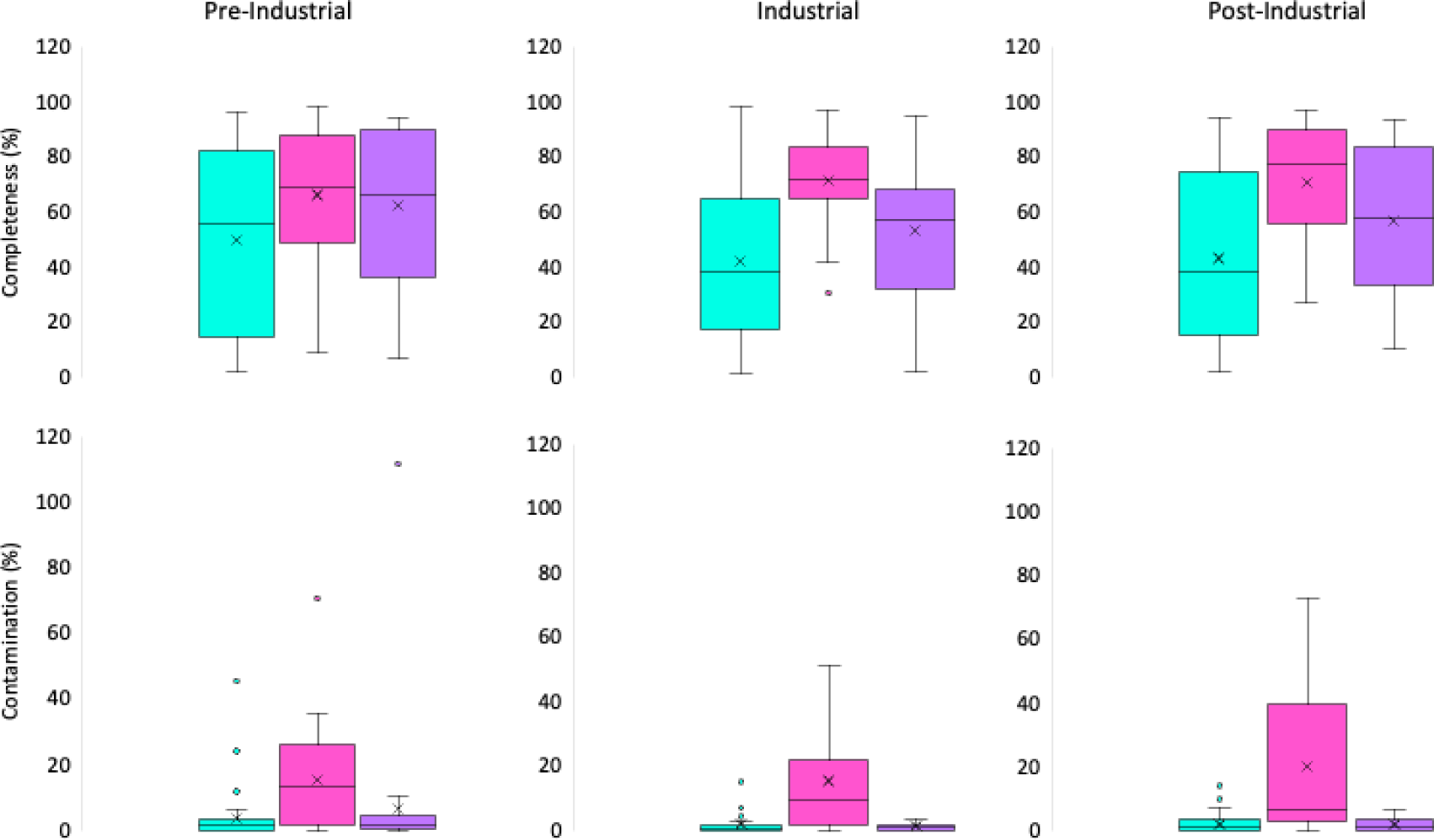
Varying contamination and completeness statistics (CheckM) for CONCOCT, Maxbin, and MetaBAT bins (before DAS Tool selection) across the Pre-Industrial, Industrial, and Post-Industrial periods.

**Table 2.**
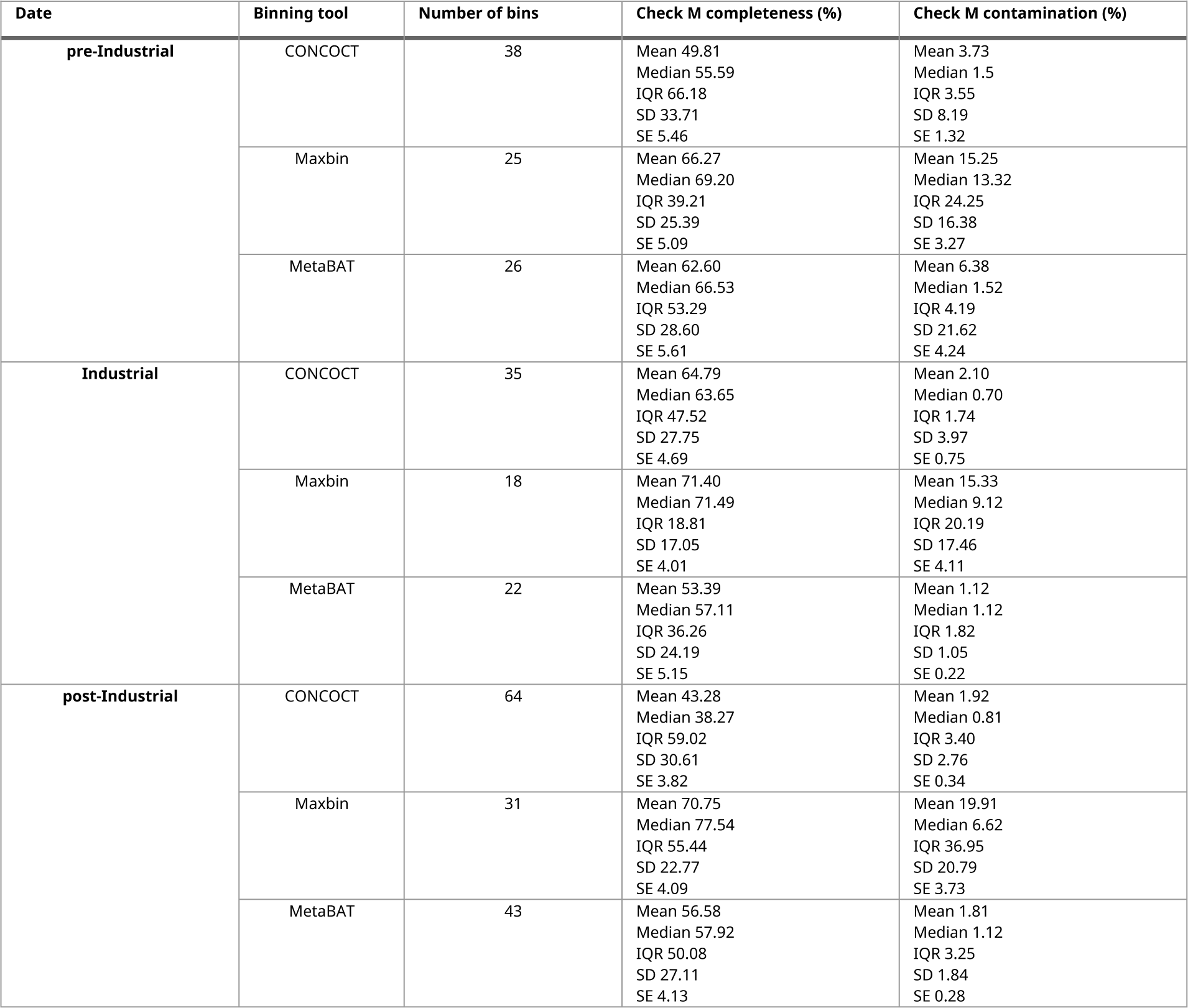
Table of Check M completeness and contamination values for bins, including bin period, binning tool, and number of bins.

## Discussion

### Modern MAG binning tools work on aDNA

To determine if MAG retrieval tools, as they are commonly integrated into pipelines for recovery of MAGs from modern microbiome samples, are also effective on aDNA, we first applied a standard pipeline (**Fig 1**) containing three binning tools (CONCOCT, MetaBAT, Maxbin) to 20 simulated modern and ancient datasets. MAG recovery success in modern simulated samples (85.5%) was greater than in ancient samples (which included the same reads, with simulated damage) (79.5%), but this difference was not statistically significant (p-value = 0.94). The pipeline did not produce false positives (absent MAGs) for either group. This suggests that under controlled conditions, i.e. with simulated data, modern binning tools can be applied to aDNA with some degree of success.

Ancient and modern metagenomics lack studies on links between OTU abundance and genome size and MAG recovery; however, this research shows that OTU abundance in a sample is important and that its combination with OTU genome size plays a key role in MAG recovery. OTUs with low sample abundance and large genome sizes have an increased chance of MAG recovery failure. While this is an issue for both groups, ancient samples have a larger risk, due to sequence degradation impacting sequence similarity. Read count and abundance did not seem to affect MAG success in any group.

Factors that impact MAG success such as abundance and genome size are likely to be complicated in ‘real’ aDNA samples due in part to different rates of DNA degradation linked to factors such as time, soil pH, and temperature, which may decrease the accuracy of MAG retrieval. Additionally, sequencing strategy may also impact MAG retrieval success in ancient samples. In our simulated data set, the read length was 300 bp. While this isn’t unreasonable for modern data with contemporary sequencing technology, DNA fragmentation linked to the factors mentioned above may make recovering fragments of this length from aDNA samples difficult. Certainly, most aDNA samples sequenced pre-industrial will not meet this read length standard. This means that one might anticipate that aDNA samples with varying degrees of fragmentation and degradation linked to time since burial and burial conditions, will likely have reduced MAG retrieval success. The older a sample (or the worse its preservation), the higher the likelihood of false negative OTUs. This may be compounded further by the sequencing read length strategy used, with short read lengths in combination with high DNA degradation likely to have a high impact on MAG retrieval success.

In addition to read lengths and damage, another potential factor is the impact of sequencing depth on MAGs retrieval. A MAGs simulation study (Steinke et al. 2024) found that even slight changes, as minimal as 1%, in coverage depth can affect the recovery of a MAG. Shallow sequencing depth may thus hinder the representation of low-abundance OTUs in MAGs. Recent pipelines combining third-generation and ultra-deep second-generation sequencing have improved sequencing depth for low-abundance subpopulations in human microbiomes and enhanced binning performance ( Jin et al. 2022). Therefore, if low genomic coverage is significantly affecting the retrieval of modern MAGs, the field must consider developing better extraction and sequencing techniques for aDNA.

### Single vs. combined binning tools on aDNA

For the real archaeological metagenomes, each binning tool produced a different number of bins with different levels of contamination and completeness (see **Fig. 3**), with Maxbin producing bins showing contamination levels as high as 19.91% while the other two produced bins below 7%, despite having used the same data. The differences in binning results raise questions about which tool is performing the best and whether a multi-binning strategy is necessary at all.

Using simulated data, we established that combining all three binning tools is likely to outperform a pipeline that uses only one of the binning tools, as the three tools found completely overlapping bin sets in only 44% of cases. This likely means that when applied to degraded ancient data, using a pipeline which employs only one binning tool is likely to lead to a significant underestimation of OTUs in any given sample. Again, given that this is simulated ancient data, with only moderate degradation and fragmentation on 300 bp paired-end data, this may be an underestimation of the number of OTU MAGs that may be missed with ‘real’ aDNA which will have variable degradation and potentially shorter read lengths.

This type of research regarding binning combination to look at OTU representation in MAGs is lacking in the field of ancient and modern metagenomics so there are not many studies to compare this to. But we do know that other research yielded the most high-quality bins when combining multiple bins on modern metagenomic data compared to using just one binning tool (Uritskiy et al. 2018; Yue et al. 2020; Rühlemann et al. 2022). More research like this using simulated data with artificial damage would be advantageous in the field of ancient metagenomics because then we would be able to see the effect that damage has on MAG success.

### Optimal amount of samples for MAG assembly

To understand factors affecting MAG retrieval success in real archaeological data, we applied our multi-binning pipeline approach to real aDNA from dental calculus samples across multiple periods using different read lengths. With a ‘single sample’ approach, in which we applied the pipeline to each sample individually, the results were relatively striking, namely, retrieving MAGs is frequently not possible. This may not be surprising, given that aDNA samples will be fragmented, often poor quality, and samples may well be sequenced from relatively low quantities of DNA. The abundance of any given OTU in the sample may also play a key role. While samples with longer read lengths had slightly higher retrieval success, it is clear that a single-sample approach is not meaningful with aDNA, and that grouping samples, based on meaningful parameters (sample age, location etc) is a more appropriate strategy for MAG retrieval. Some of these samples are likely part responsible for GTDB-Tk not always being able to assign family, genus, or species of bacteria to MAGs, and it would be useful if more studies were to investigate the binning of different periods to further investigate how read count/damage (‘ancientness’) of certain samples contributes to poorly defined MAGs (taxonomically undeterminable past family level).

## Conclusion

This research has revealed that modern MAGs binning tools can retrieve medium to high quality MAGs that reflect original OTUs from highly damaged reads. Our findings have also revealed that a combined binning approach on multiple samples is more beneficial than binning single samples and using one tool to bin samples. We emphasise the importance of testing the suitability of modern tools on simulated data in ancient metagenomics as aDNA has varying characteristics that may affect MAG success. Moreover, we urge for more research to investigate the impact of damaged aDNA with low-coverage short-read length on MAG creation.

## Acknowledgments

The authors acknowledge the use of the University of Bradford High Performance Computing Service in the completion of this work.

## Author Contributions

Conceptualization: A.T., C.J.M. Supervision: A.T., C.J.M., and C.S. Funding acquisition: C.S., A.T., C.J.M. Investigation: F.J.S, G D-A., and A.T. Formal Analysis: F.J.S, G. D-A., A.T. and C.J.M. Data Curation, Visualization and Writing -Original Draft: F.J.S. Writing - Review & Editing: all authors.

